# A Multimodal Workflow for Spatial Metabolic Neighborhood Mapping in Neural Rosette Cultures

**DOI:** 10.64898/2026.04.13.715964

**Authors:** Oluwatomisin N. Adebayo, Akhil Turaga, Minjae Chung, Facundo M. Fernández, Melissa L. Kemp

## Abstract

Neural rosettes are hallmarks of the neural progenitor cell stage that is a necessary pre-condition for manufacturing central nervous system lineages. Characterization of early changes during differentiation through positional arrangement and metabolic shifts that occur in a multi-day protocol would facilitate cell culture quality monitoring and optimization of batch culture yield. We describe an analytical framework for identifying neural rosettes from confocal microscopy within a colony of differentiating stem cells and translating co-registered, cell-resolved MALDI imaging data into interpretable readouts that are compatible with cell manufacturing needs. Rather than evaluating hundreds of ion images sequentially, the pipeline converts each region of interest into a single-cell feature matrix and summarizes whole-spectrum variation using PCA, graph-based Leiden clustering, and UMAP visualization. The resultant metabolic neighborhoods provide quantification of molecular heterogeneity within colonies and - when mapped back to x–y space - form coherent spatial domains. Together, these outputs create a practical bridge between multimodal MALDI capabilities and process-relevant interpretation: neighborhoods can be compared across conditions, ranked markers can be prioritized as putative critical quality attributes, and spatial organization can be quantified without manual, feature-by-feature screening.

## Introduction

Self-organization of neural rosettes, radial arrangements in differentiating ESC or iPSC culture, mimics features of embryonic neural tube formation and neuroectoderm specification. Neural rosettes are a signature of neuroprogenitors, capable of further differentiation into neurons, oligodendrocytes, and astrocytes; the in vitro manufacturing of neural lineages derived from pluripotent populations relies upon successful achievement of this stage as a hallmark of production (Elkabetz et al., 2008; Wilson & Stice, 2006). Recent efforts to regularize neural rosette formation through surface micropatterning(Aghayee & Ashton, 2021; Knight et al., 2018; Lundin et al., 2024) are promising for applications of in vitro toxicological screening and studies of multicellular coordination associated with neurulation, yet many needs for neural rosette monitoring (such as quality control of neural progenitor cells) require detection and characterization within a colony or organoid tissue without geometric constraints. Prior computational tools developed for analyzing neural rosette properties have focused on longitudinal tracking to use cellular speed and angular motion to compare features of early radial glial rosettes versus mid radial glial rosettes rather than identifying initiation of lumen formation(Ziv et al., 2015).

Metabolic imaging has demonstrated key advantages for monitoring loss of pluripotency and lineage specification in stem cell colonies. Techniques such as matrix-assisted laser desorption/ionization (MALDI) and desorption electrospray ionization (DESI) mass spectrometry imaging (MSI) have improved resolution to routinely image at 5-50 μm and are capable of detecting hundreds of metabolic features per pixel, yielding very high-dimensional spatial information of differentiating cells; for example, using co-registration with bright field(Priyadarshani et al., 2024) or confocal microscopy yields near single cell information(Nikitina et al., 2020). In addition, metabolic changes occur during loss of pluripotency much earlier than canonical transcription factor changes such as Oct4, delivering new insight into metabolic priming hours to days before expression changes are evident(Nikitina et al., 2023). Finally, the response of cellular metabolism to subtle changes in media composition and culture reagents provides sensitive metrics of quality control during cell differentiation associated with manufacturing processes.

Here, we present a multimodal analysis workflow that converts live cell microscopy and high-dimensional MALDI spectra into spatially interpretable, holistic metabolic maps to evaluate spatial metabolomic features associated with neural rosette organization. The combined computational/experimental pipeline interprets a coordinate-resolved matrix (x,y positions with m/z intensities) and outputs reconstructed metabolic neighborhoods associated with rosette membership. We frame how these metabolic clusters can be interpreted alongside phenotypic surface markers and extended toward early rosette-relevant cues by integrating automated rosette detection and complementary spatial statistics for center-to-edge enrichment mapping. This pipeline can be deployed across conditions to accelerate reagent/perturbation screening and improve the interpretability and standardization of MSI analysis of differentiating cells.

## Cell Maintenance & Differentiation

Human embryonic stem cells (WA09 “H9”; WiCell) were maintained in 6-well plates for three passages prior to differentiation on plates coated with 1 mL growth factor–reduced Matrigel (1:100 in KnockOut DMEM) per well. Cells were fed daily with 2 mL per well of maintenance medium (mTeSR basal medium plus supplement, 4:1) until the first passage. For passaging, cells were incubated with 0.5 mL Accutase per well at 37 °C for 5 min, gently lifted, and transferred to a 15 mL conical tube containing ∼5× the Accutase volume of medium. Cells were centrifuged at 0.2 rcf for 5 min, the supernatant was removed, and the pellet was resuspended in 2 mL E8 medium (E8 basal medium plus supplement) containing 2 µL ROCK inhibitor for cell counting. Cells were then reseeded at 150,000 cells per well in 2 mL E8 medium with 2 µL ROCK inhibitor for the first 16–18 h; thereafter, medium was replaced with E8 medium without ROCK inhibitor until the next passage. After two passages in E8, cells were transitioned into the neural differentiation workflow.

For neural rosette differentiation, cells were dissociated again using 0.5 mL Accutase per well (37 °C, 5 min), collected into a 15 mL tube with ∼5× medium, and centrifuged at 0.2 rcf for 5 min. The pellet was resuspended in 2 mL E8 medium supplemented with 2 µL ROCK inhibitor and reseeded at 175,000 cells/cm² (e.g., 1,680,000 cells per 6-well) in 4 mL E8 medium containing 4 µL ROCK inhibitor for the first 16–18 h. Medium was then replaced with E6 medium (Gibco) and cultures were maintained in E6 with daily medium changes for 8 days to promote rosette formation.

## Live Cell Microscopy

After 8 days of differentiation, cultures were stained live prior to MALDI imaging. Medium was aspirated and cells were gently washed three times with PBS (2 min per wash). Live-cell antibodies NL557-conjugated Mouse Anti-Human SSEA-1, and Alexa Fluor 647-conjugated Mouse Anti-Human NCAM-1/CD56 were diluted 1:50 each in pre-warmed E6 medium and applied to the wells. Cultures were incubated with the antibody cocktail for 45 min, after which the staining solution was aspirated, and cells were gently washed once with PBS. Fresh E6 medium was then added, and samples were immediately transferred for imaging on a Nikon W1 spinning-disk confocal microscope using a 10× objective (0.65 µm/pixel). Imaging was performed using protocol described in (Nikitina et al., 2020). A live-cell chamber at 37 °C and 5% CO₂ with identical acquisition settings across conditions to visualize live-cell staining. For each well, at least two non-overlapping fields of view were acquired on a single focal plane. Images were saved in ND2 format and processed using FIJI for downstream analysis.

## Automated rosette detection for ROI

Following live-cell microscopy acquisition, automated detection of neural rosette structures was performed to define regions of interest (ROIs) for downstream MALDI-MSI analysis. The detection pipeline was developed to identify rosette candidates based on characteristic features of spatial distribution patterns (contrast, relative area, circularity) observed in DAPI nuclear staining, without requiring geometric constraints or micropatterning.

DAPI-stained confocal microscopy images (ND2 format) were first converted to TIFF format. Adaptive local contrast enhancement (CLAHE; clip limit = 3.0) was applied to enhance intensity features and improve discrimination of rosette-associated nuclear density patterns from background regions, then subjected to Gaussian Blur smoothing (σ=15-19) to reduce noise and smooth intensity gradients to enhance rosette boundary while preserving overall structural features.

Rosette centers, characterized by chromatin-depleted lumen regions, were identified by intensity thresholding on the blurred image. Pixels below the 10th percentile of intensity were classified as dark regions and masked accordingly. Connected component labeling was applied to identify spatially contiguous dark regions, with each labeled region representing a candidate rosette structure. A percentile-based size filter was applied to the detected dark regions for removal of imaging artifacts, noise, and abnormally sized regions. Regions were ranked by area, and those falling below the 80th percentile (small noise artifacts) or above the 95th percentile (oversized artifacts or edge effects) were excluded. This percentile-based approach proved more robust than absolute size thresholds across varying imaging conditions, preserving biologically relevant rosette structures while systematically removing technical artifacts.

To prioritize the most confident detections, a weighted scoring system was implemented combining normalized size (70%) and circularity (30%) metrics. The 70% weighting on size reflects the biological significance of rosette lumen prominence, as larger dark regions indicate more defined rosette structures with clear centers. The 30% weighting on circularity captures the radial symmetry characteristic of neural rosette organization around the central lumen. Both metrics were normalized to a 0-1 scale to enable fair comparison across the size range, and candidates were ranked by their composite weighted score. The top 20 ranked rosettes were mapped back onto the original CLAHE-enhanced DAPI image with color-coded numerical labels indicating rank. Detected ROI coordinates were exported as a table containing rosette ID, centroid position, area, circularity, weighted score, and geometric metrics.

## Sublimation

Samples were briefly washed to reduce residual salts prior to MALDI-MS acquisition by immersing them in 5 mM ammonium formate for 3s, then immediately air dried prior to matrix deposition. For external calibration, 1 µL of red phosphorus was spotted adjacent to the sample area.

Norharmane was deposited by sublimation, processing one slide at a time at 250 °C under vacuum (1.1 × 10⁻¹ Torr) for 6 min to produce a uniform matrix coating suitable for MALDI.

## MALDI Acquisition

Matrix-coated samples were analyzed in reflectron mode on a RapifleX TissueTyper time-of-flight (TOF) mass spectrometer (Bruker Daltonics, Billerica, MA, USA) equipped with a Smartbeam3D 10 kHz Nd:YAG laser (355 nm). Imaging acquisition was controlled using FlexImaging 4.0 (Bruker Daltonics) with the Smartbeam3D single-beam setting and a raster step size (pixel size) of 5 µm in both x and y. Spectra were acquired in negative ion mode over m/z 400–1600, averaging 200 laser shots per pixel. External mass calibration was performed using red phosphorus prior to data acquisition to ensure accurate m/z assignment.

## Co-registration

MALDI-MS imaging data were preprocessed in SCiLS Lab (SCiLS GmbH, Bremen, Germany) during data import, including baseline correction and total ion count (TIC) normalization. Peaks were defined using a fixed tolerance window (5 mDa) around the mean peak center to account for small drifts. Peak detection was performed in SCiLS Lab and refined by selecting m/z features that exhibited signal localized to the cell colony region. To enable downstream multimodal analysis, a Python-based export workflow was used to extract ion intensities for the selected m/z features from raw rapifleX data and generate corresponding ion images for registration.

Co-registration between confocal microscopy and MALDI-MSI was performed as previously described (Nikitina et. Al, 2020) using an image alignment approach based on mutual information, treating the DAPI nuclear channel as the reference image and a composite MALDI ion image (generated by averaging selected peaks) as the moving image. Cells were segmented from the confocal images by first identifying nuclei and then expanding these regions to define individual cells. Following segmentation, each cell mask was overlaid onto the registered MALDI ion images to quantify per-cell m/z intensities; corresponding per-cell fluorescence intensities were also extracted from the confocal channels. The resulting output was a single cell–resolved table containing cell coordinates/shape descriptors and matched MALDI and fluorescence intensities, exported as a CSV for downstream dimensionality reduction and clustering.

## Preprocessing

After compiling per-cell intensities into CSV format, MALDI intensities were total-ion normalized per cell so that ion abundances were comparable across cells, and values were log-transformed to stabilize variance and reduce the influence of very large peaks. Rule-based isotopic filtering was applied to the exported m/z columns prior to normalization and clustering assuming singly charged ions (z =1) in negative mode ([M–H]⁻). For each m/z feature, companion peaks were searched at +1.0033548 Da and +2.0067096 Da, the normal ^13^C isotope spacing for singly charged ions, using a ±0.01 Da matching window to accommodate small peak shifts. When matches were identified, the higher-mass isotope columns (M+1 and M+2) were removed, and the monoisotopic column was retained. The resulting filtered m/z was used as input for downstream clustering and UMAP visualization.

Principal component analysis (PCA) on m/z features was performed, retaining the top 20 principal components for downstream analysis. A k-nearest neighbor (KNN) graph (k = 15) was subsequently constructed in PCA space to represent local metabolic similarity between cells, and Leiden community detection (resolution = 0.6) was applied to partition the graph into clusters.

This workflow assigns a cluster label to each cell and reveals distinct metabolic neighborhoods within the region of interest, which were subsequently visualized in a UMAP embedding.

## UMAP

A UMAP embedding was then computed from the PCA representation to visualize relationships among cells in a low-dimensional space while preserving local neighborhood structure.

Differential feature ranking was performed on the m/z intensity matrix using Scanpy’s rank_genes_groups framework, treating each Leiden cluster as a group and all remaining cells as the reference. For each cluster, candidate markers were ranked by effect size (log fold change) and filtered for significance (p-value < 0.05). The top five m/z features per cluster were retained as cluster-enriched markers. In UMAP plots, each point represents a single cell, and its color reflects the abundance of the top m/z value or cluster label. In addition, cluster assignments were mapped back into the original coordinate space (x,y) to generate spatial cluster maps and confirm that UMAP-defined metabolic neighborhoods corresponded to coherent spatial domains in the sample.

## Radial Gradient Analysis

Post co-registration of MALDI-MSI with fluorescence microscopy, single cell coordinates with assigned rosette identifiers were extracted from the segmentation and co-registration pipeline output of the raw MALDI imaging data. Per-cell coordinates with neural rosette labeling from the rosette detection pipeline were merged to the coordinate table using matched spatial positions (x, y). For each rosette, cells were partitioned into “Lumen” and “Non-lumen” populations based on their relative position within the structure: cells within the innermost 20th percentile of distances from the geometric centroid were labeled as “Lumen”, with the remaining cells designated as “Non-lumen”. This partitioning established an internal reference boundary for subsequent radial measurements. For each cell, the Euclidean distance to the nearest Lumen-labeled cell was computed using a k-dimensional tree (KDTree) data structure, yielding a radial coordinate *r*. This distance was then normalized independently within each rosette as 𝑟*_𝑛orm_*= 𝑟/ max(𝑟), scaling radial positions to a common [0, 1] interval regardless of absolute rosette size or shape. This normalization enables cross-rosette comparisons by mapping all cells onto a common radial axis where 0 represents the lumen boundary and 1 represents the outermost periphery.

## Radial Binning and Gradient Quantification

Cells were assigned to discrete radial bins based on their normalized distance 𝑟*_𝑛orm_*. A fixed 8-bin quantile scheme was used as the primary analysis method, wherein cells were partitioned into five groups of approximately equal size using quantile-based boundaries on the rank-transformed 𝑟*_𝑛orm_* values. This approach ensures balanced statistical power across bins regardless of cell density gradients. Each bin was assigned a normalized radial position (0 = lumen, 1 = periphery) computed as the bin midpoint, enabling Spearman correlation between metabolite intensity and radial position. For each m/z feature, the change in fractional abundance relative to the lumen was computed by a delta value Δ log_2_ 𝐼 = log_2_(𝐼_𝑖𝑖_) − log_2_ (*Ī_lumen_*), representing the difference between the mean fractional abundance in each radial bin and that in the innermost (lumen) bin.

### Correlation and Permutation Analysis

Spearman rank correlation coefficients (ρ) were computed between the delta values and radial bin positions for each m/z feature within each rosette. Next, a curated set of candidate rosette-enriched m/z features (“GOOD” set), identified from prior UMAP clustering as features with elevated signal in rosette-center-associated clusters, was compared against a null distribution by permutation testing. A comparison set (“BAD” set) established a baseline for non-gradient behavior. While the permutation test typically utilizes a randomly sampled null set, the “BAD” subsets displayed in Figure 5-b, c, d were specifically selected for visualization purposes to represent features with no prior information. For each 200 permutation iterations, a random subset was drawn without replacement and mean Spearman correlation was computed. The empirical p-value was calculated as the proportion of random samples achieving a mean correlation equal to or greater than the observed value for the curated set. This procedure tests whether the selected m/z features exhibit stronger radial gradients than expected by chance.

## Results & Discussion

### Establishing neural rosette cultures for single-cell MALDI–UMAP analysis

Embryonic stem cell–derived neural rosettes were generated from WA09 (H9) colonies cultured on indium tin oxide (ITO) slides and differentiated using an Essential 6 (E6)-based neural induction framework, with rosette-like neuroepithelial organization apparent by day 8. After differentiation, colonies were live-stained and imaged to confirm rosette-associated phenotype in situ and generate the fluorescence reference images used for downstream multimodal alignment (Figure 1). Because MALDI acquisition and fixation-based immunostaining are not compatible on the same sample, parallel slides were processed in tandem: one slide was used for MALDI-MSI, while a matched slide was fixed and stained to validate rosette structure.

**Figure 1.**
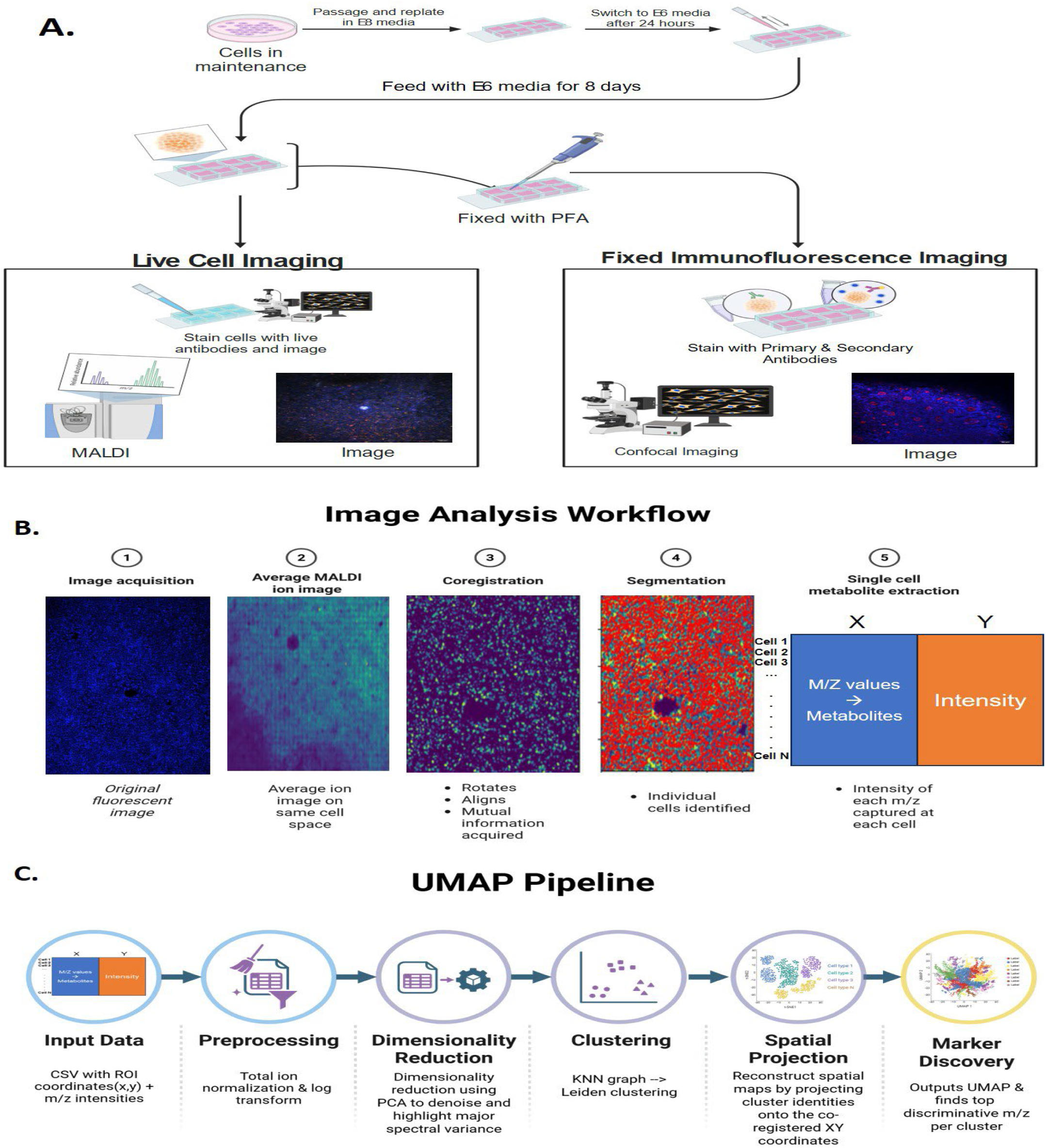
Identification of rosettes in H9 colonies and an overview of the MALDI–UMAP analysis workflow. (A) Schematic of the neural differentiation timeline on ITO slides (maintenance → E8 attachment phase → E6 differentiation through day 8). Representative experimental design of live-cell fluorescence images of WA09 (H9) colonies on day 8 live stained to confirm rosette-associated phenotype and define rosette-containing regions for downstream analysis. Parallel slides were processed in tandem with identical culture conditions and live staining; one slide proceeded to MALDI-MSI acquisition, while the matched slide was fixed and immunostained to validate rosette epithelial organization. (B) Multimodal co-registration of MALDI ion images to fluorescence microscopy, enabling extraction of per-cell m/z intensities using cell segmentation masks and aligned coordinates. (C) Data reduction and visualization workflow: aligned single-cell MALDI intensities were compiled into a CSV matrix (cell coordinates × m/z features), followed by normalization, PCA, k-nearest neighbor graph construction, Leiden clustering, and UMAP embedding to reveal metabolic neighborhoods within the region of interest.

Because the size of the imaged region is a rate-limiting step in MALDI-MSI, we first developed a pipeline for detecting rosettes from DAPI-stained cells. Visual inspection of detected rosette candidates overlaid on fluorescence images confirmed correspondence with rosette-like morphology (Figure 2). Validation against ground-truth rosette annotations (identified by high ZO-1 expression) demonstrated that the model successfully detected biologically relevant rosettes as the ‘top 10 ranked candidates’, indicating strong precision in prioritizing high-confidence detections. This automated approach enabled scalable, reproducible ROI selection across multiple fields of view and experimental conditions for downstream multimodal analysis.

**Figure 2.**
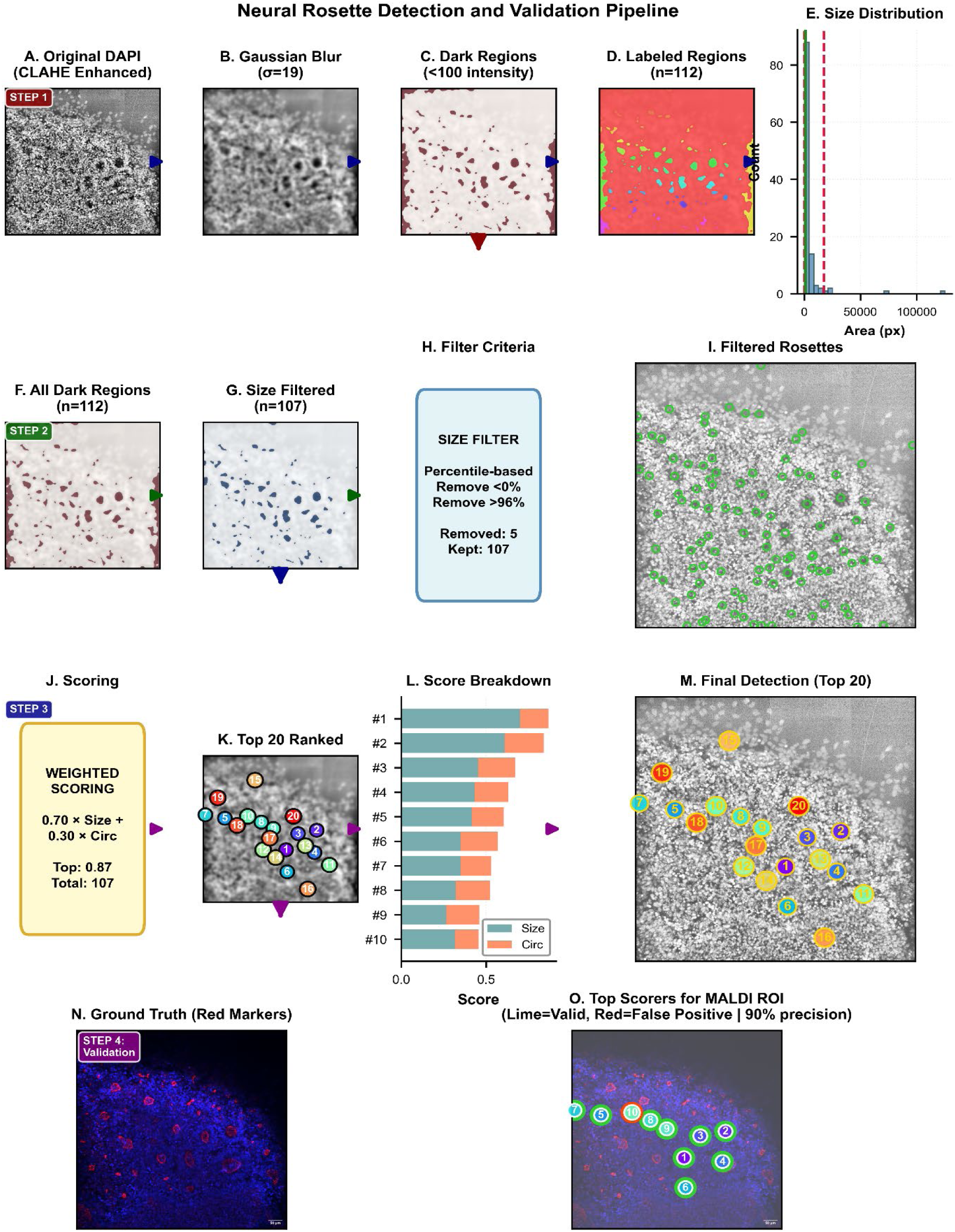
Automated Neural Rosette Detection Pipeline from DAPI Imaging to Validated Candidate Selection. (A) DAPI-stained nuclei were enhanced using adaptive histogram equalization (CLAHE; clip limit=3.0) to improve local contrast and reveal nuclear organization patterns. (B) Gaussian smoothing (σ=19) was applied to reduce noise and enhance rosette boundary visibility as continuous dark regions while preserving structural features. (C) Pixels below the 10th percentile of intensity were classified as dark regions corresponding to chromatin-depleted rosette lumen centers. (D) Connected component labeling identified spatially contiguous candidate regions for downstream filtering. (E) Histogram of detected region areas showing heterogeneous size distribution, with red dashed lines indicating percentile-based thresholds (80th and 95th percentiles) used for outlier removal. (F) All detected dark regions before size filtering, including imaging artifacts and noise. (G) Regions above the 96th percentile (oversized artifacts) were systematically removed, retaining biologically relevant candidates. (H) Percentile-based filtering criteria summary showing removal of regions and retention of candidates based on size distribution. (I) Size-filtered candidates overlaid on original CLAHE-enhanced DAPI image with lime circles confirming spatial correspondence with visible chromatin-depleted lumen structures. (J) Weighted scoring methodology combining normalized size (70% weight, reflecting lumen prominence) and circularity (30% weight, capturing radial symmetry), yielding top score from total ranked candidates. (K) Top 20 ranked detections color-coded by rank on blurred DAPI image with numbered labels indicating rank order. (L) Score decomposition via stacked horizontal bars showing relative contributions of size (blue, 70%) and circularity (coral, 30%) components for top 10 candidates, validating weighting strategy effectiveness. (M) Final detected rosettes with top 20 ranked candidates mapped onto original CLAHE image using color-coded numbered circles. (N) Ground truth validation showing RGB image with manually annotated rosettes marked in red for accuracy assessment. (O) Top scorers for MALDI ROI: validation overlay with top 10 detected candidates on ground truth image, where lime circles indicate true positive detections (overlap with red markers) and red circles indicate false positives (detection radius: 100 pixels).

MALDI-MSI was performed to map metabolite and lipid signals across the same colony regions at high spatial resolution, allowing detection of molecular signals that localize to rosette regions (Figure 3). Multimodal co-registration was performed to align MALDI ion images to the corresponding fluorescence images and enable single-cell resolved extraction of MALDI intensity values using an established coregistration workflow to extract per-cell features (Nikitina, JASMS, 2020). The aligned data were converted into a cell-resolved matrix where cell coordinates are matched with individual m/z intensities, which serves as the direct input to the UMAP workflow.

**Figure 3:**
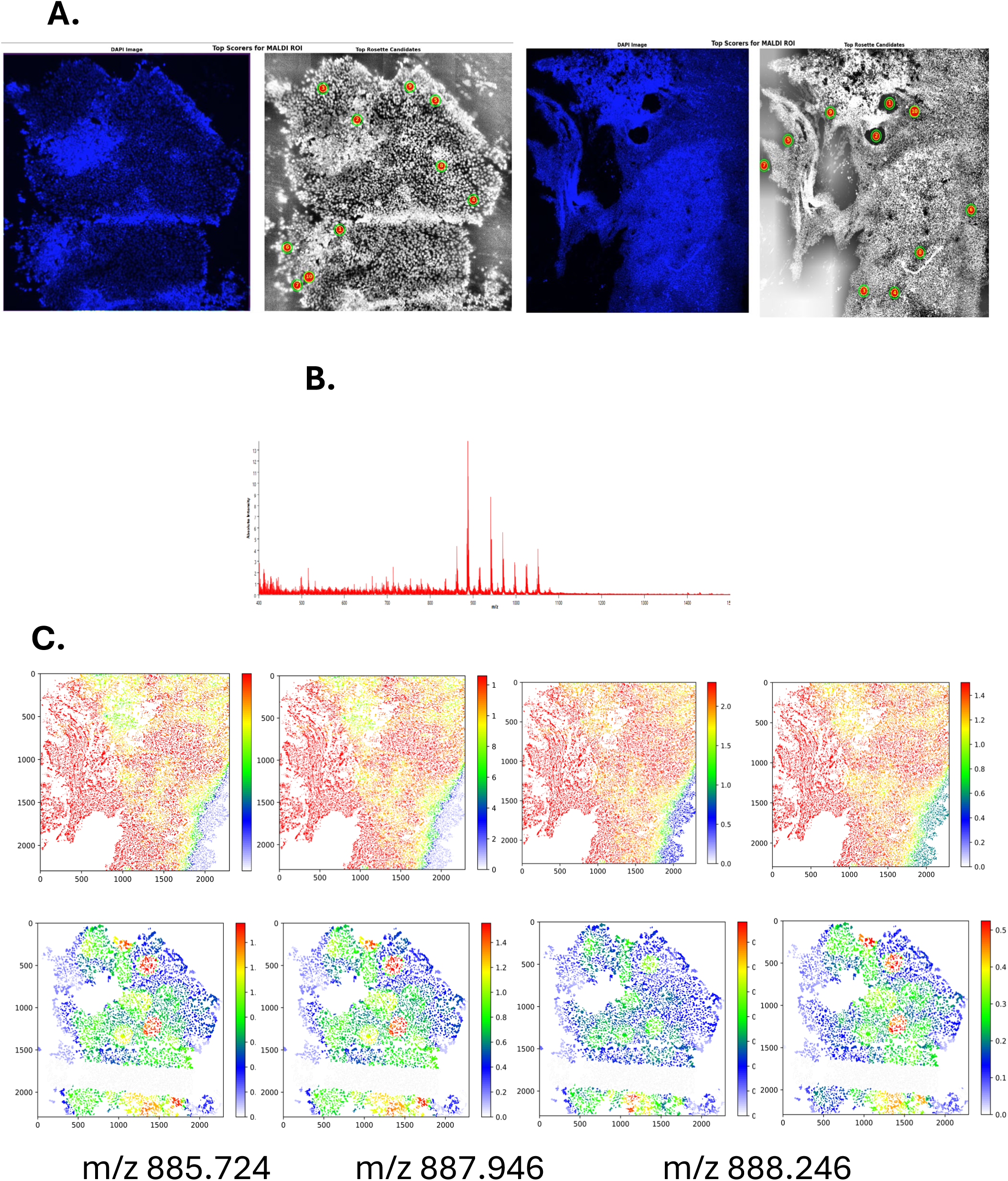
MALDI-MSI identifies rosette-enriched metabolic features. (A) DAPI live cell image being put through the automatic rosette detector to identify putative rosette ROIs for MALDI(B) Average negative-ion MALDI spectrum collected from the imaged colony region (m/z 400–1600). (C) Ion images for m/z features enriched in rosette-containing regions, including m/z 885.724, 887.946, 888.246, and 914.741, shown as spatial variance maps over the same field of view.

Rather than going through hundreds of ion images individually, the resulting pipeline summarizes the full spectral landscape into metabolic neighborhoods through graph-based clustering and UMAP visualization, providing an interpretable map of how metabolic states are organized within the region of interest.

## MALDI-MSI identifies rosette-enriched metabolic features

MALDI-MSI acquisition on day 8 H9 colonies produced robust negative-ion spectra across the imaging range (m/z 400–1600), yielding a dense set of molecular features for spatial interrogation. Inspection of individual ion images revealed pronounced spatial heterogeneity within the colony, with multiple lipid-associated signals showing localized enrichment in rosette-containing regions rather than uniform distribution across the field. Several m/z features were consistently elevated in rosette-rich areas, including m/z 885.724, 887.946, 888.246, and 914.741 (Figure 3). These ions formed spatially coherent hotspots that aligned with rosette-like morphology observed by live-cell imaging, supporting the presence of a distinct metabolic program associated with rosette organization.

Although these representative ion maps highlight candidate rosette-enriched features, interpreting MALDI data peak-by-peak does not scale. Each dataset contains hundreds of informative m/z variables, many of which covary or capture overlapping biology. This motivated our workflow for systematic identification of cluster-defining m/z markers rather than manual inspection of individual ion images.

## PCA, Leiden clustering, and UMAP reveal spatial metabolic neighborhoods

After total-ion normalization and log transformation, principal component analysis (PCA) was applied to denoise and compress the high-dimensional m/z space into a lower-dimensional representation while preserving dominant sources of biological variation. The top 20 principal components were retained for downstream analysis. In this PCA space, cells with similar metabolic profiles cluster closer together, providing an initial indication that the colony contains structured metabolic co-variation.

To define this structure objectively, a k-nearest neighbor (KNN) graph was constructed in PCA space so that each cell was connected to its most metabolically similar neighbors, capturing local relationships in the full multivariate spectrum rather than relying on any single m/z feature. Leiden community detection was then applied to this graph to assign a discrete cluster label to each cell, partitioning the region of interest into metabolic neighborhoods with shared spectral signatures (Figure 4). UMAP was subsequently computed from the same PCA representation to visualize these relationships in two dimensions, placing cells with similar spectral profiles close together while separating cells with distinct profiles. Viewing the marker intensity map next to the cluster map facilitates assessment of cluster-enrichment of a given metabolite. This provides an intuitive check that the top-ranked m/z features are specific to the metabolic neighborhood they are intended to define, and it highlights gradients or substructure within and between clusters when present.

**Figure 4:**
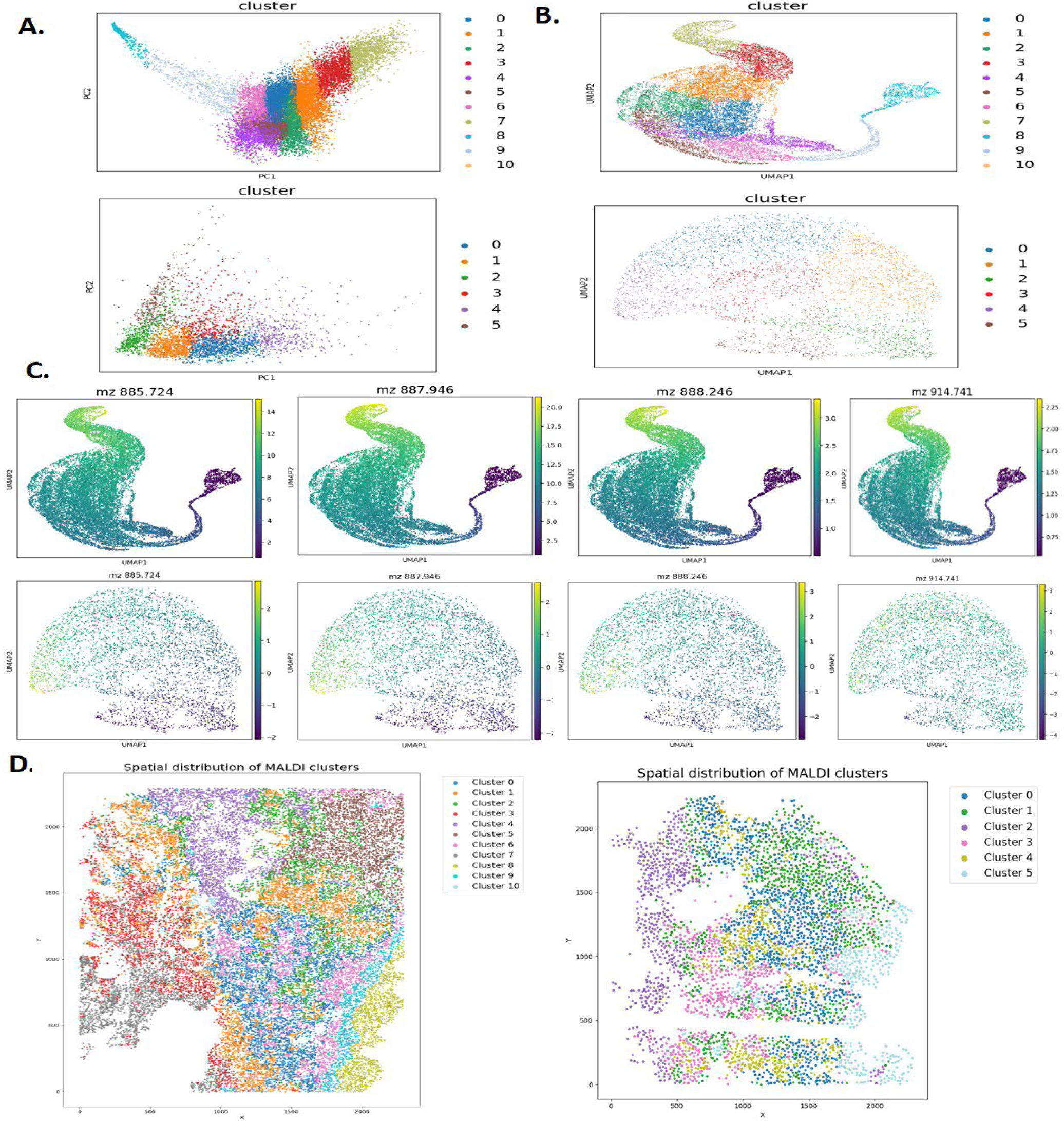
PCA, Leiden clustering and UMAP. (A) Principal component analysis (PCA) of the cell-resolved MALDI feature matrix (log1p, TIC-normalized) colored by Leiden cluster identity, showing separation of metabolic states in PC space (PC1 vs PC2). (B) UMAP embedding computed from the PCA representation, colored by Leiden clusters, visualizing relationships among metabolic neighborhoods. (C) UMAP feature overlays for selected cluster-enriched ions (m/z 885.724, 887.946, 888.246, and 914.741), demonstrating that top-ranked m/z features localize to specific regions of the embedding and track neighborhood structure. (D) Spatial back-projection of Leiden cluster labels onto the original x–y coordinates (tissue space) reveals that metabolic neighborhoods correspond to coherent spatial domains within the colony

To connect the embedding back to colony architecture, Leiden cluster identities were projected onto the original x,y coordinates used for MALDI acquisition and microscopy alignment. Back-projection revealed that clusters formed organized spatial domains rather than random mixtures of labels, supporting the interpretation that the computational neighborhoods reflect structured metabolic patterning within the colony. Together, PCA-driven denoising, graph-based Leiden clustering, and UMAP visualization provide a scalable alternative to sequential single-ion interrogation by summarizing whole-spectrum MALDI data into spatially grounded metabolic neighborhoods that can be used downstream for marker ranking and phenotype linkage.

## Radial gradient analysis reveals lumen-enriched metabolite signatures within neural rosettes

To quantify spatial patterns within the asymmetric rosette architecture, we applied a distance-to-boundary radial analysis. Neural rosettes exhibit irregular, often elongated morphologies that deviate substantially from ideal circular (mean aspect ratio > 1.0; Figure S2), enclosing a central lumen with an irregular boundary that serves as the architectural anchor for radial organization(Filan et al., 2024),(Knight et al., 2018).To accommodate this shape variability, a distance-to-boundary model was employed rather than a conventional distance-from-center approach.

Among [16] features examined via Spearman rank correlation, a subset of [8] features associated with rosette-center UMAP clusters displayed uniformly negative correlations (mean ⍴ = -0.366), indicating progressive depletion from the lumen toward the periphery (Figure 5B,D).

**Figure 5.**
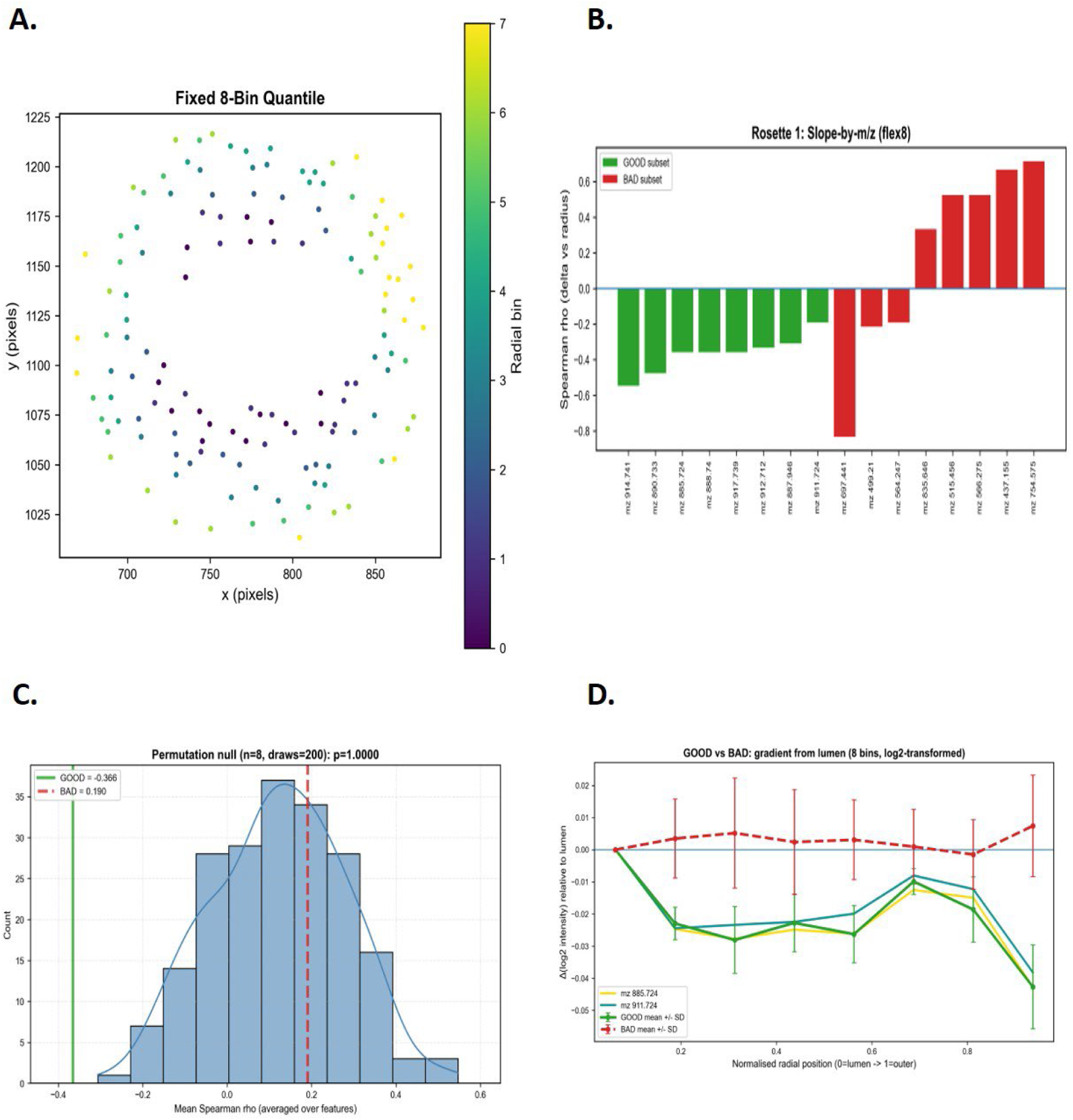
Radial gradient profiling of metabolite distributions within neural rosettes. (A) Spatial distribution of cells within a representative rosette partitioned into eight radial bins by fixed quantile binning, color-coded by bin assignment from lumen (bin 0) to periphery (bin 7). (B) Log-transformed intensity profiles relative to the lumen bin (delta) for the lumen-enriched subset (green, solid) and comparison subset (red, dashed) plotted against normalized radial position. (C) Permutation null distribution of mean Spearman ρ from [200] random draws of 8 m/z values(histogram), compared with the observed mean for the lumen-enriched subset (solid green vertical line, ρ = [−0.366]) and comparison subset (dashed red vertical line, ρ = [0.190]). (D) Spearman rank correlation coefficients (ρ, delta vs. radial position) grouped by lumen-enriched (green) and comparison (red) subsets. Annotated m/z features (**885.724: yellow**, **911.724: teal**) represent known iPSC differentiation predictors overlapping with lumen-enriched (’GOOD’) subsets. Negative ρ values indicate lumen enrichment; positive values indicate peripheral enrichment.

Representative features, including *m/z* 914.741, exhibited monotonic declines across all radial bins. (Figure. S1-a) In contrast, features unassociated with rosette centers like m/z 956.74 exhibited spatially invariant profiles (Figure. S1-b).

To assess the statistical significance of this pattern, we performed permutation testing on the representative rosette structure. Observed lumen-enrichment was significantly distinct from random sampling; the experimental mean fell entirely outside the null distribution (n=200, random draws), which exhibited a slight edge-enrichment bias of ⍴ ∼ +0.2 (Figure 5D).

Several lumen-enriched features are consistent with phosphatidylinositol (PI) species. Notably, PI 38:4 (m/z 885.6) and PI 40:5 (m/z 911.5, shown in Figure. 5d) were previously identified as key lipid predictors of iPSC differentiation stage, with regulation via PI 3-kinase (PI3K) signaling influencing spatial colony organization (Nikitina et al., 2023). While these results establish a quantitative link between spatial lipid gradients and established metabolic signatures of neural induction, a deeper layer of analysis using tandem mass spectrometry (MS/MS) is required to formally annotate these features. Such molecular resolution would allow interpreting these contextual signals interrogate the mechanistic pathways driving this spatial patterning phenomenon.

A crucial step toward manufacturing translation is standardizing how rosette-containing regions are defined and analyzed. Manual rosette annotation is slow and introduces user-dependent bias, limiting reproducibility across batches and perturbations. Incorporating rosette detection as an upstream layer addresses this gap by enabling consistent identification of rosette-like structures and producing reproducible ROIs for downstream MALDI analysis. This step saves instrumentation time and promotes automation where detected ROIs can be batch-processed through the same preprocessing, embedding, and marker-ranking steps, enabling direct comparisons across lots, genetic backgrounds, or targeted perturbations. This workflow gives way for a scalable assay to ensure quality readouts at one day or tracked over time.

Several alternative analysis approaches were explored to address the same core challenge: extracting reproducible, biologically meaningful information from high-dimensional MALDI images, but were limited by scalability or interpretability. Manual screening of individual ion images can identify visually striking features, yet it does not readily reveal multivariate patterns shared across many peaks and is difficult to standardize across users and experiments. ROI-averaged spectra and bulk summary statistics offer a concise readout, but they mask heterogeneity within the ROI and can wash out localized rosette-associated signals, especially when rosette and non-rosette areas are analyzed together. Single-marker cutoffs can be unreliable because signal strength varies between runs, and one m/z feature rarely captures an entire cell state. In contrast, neighborhood-based embedding uses many ions at once at single-cell resolution and produces clear outputs: cluster labels, ranked marker m/z features, and cluster maps projected back onto the sample.

With the development of this pipeline, there are limitations that motivate the next steps. Marker ions can be ranked and mapped, but confirming their molecular identities will require follow-up validation. MALDI signals can vary with matrix coating and acquisition conditions, and consistent cross-run calibration and QC samples are still needed for long-term comparisons. Co-registration and segmentation can introduce errors, especially at boundaries and in regions with strong spatial gradients, so improving masks and testing how sensitive gradient results are to segmentation choices will strengthen robustness.

In future work, the most direct extension is to integrate time-lapse microscopy with scheduled MALDI sampling to add dynamic information about the timing of rosette emergence and maturation. A time-resolved version of this framework could quantify when neighborhood shifts first appear relative to morphological rosette detection, and whether lumen-enriched gradients emerge before clear rosette structures are visible. This would enable predictive models that classify cultures based on early metabolic neighborhood signatures and forecast downstream rosette yield or organization.

## Conclusion

In summary, a manufacturing-oriented spatial metabolomics workflow was developed to support objective, scalable assessment of neural rosette cultures. Multimodal co-registration enabled extraction of cell-resolved MALDI intensities, which was analyzed to define metabolic neighborhoods within regions of interest. This approach shifts interpretation from sequential inspection of individual ion images toward structured neighborhood discovery and systematic marker prioritization. Spatial back-projection confirms that neighborhoods correspond to coherent domains within the colony. Radial gradient analysis provides an additional layer of interpretability by quantifying center-to-edge localization of candidate features within organized structures. Overall, the framework supports automated comparison of protocol variables relevant to cell manufacturing and sets the stage for longitudinal validation to determine whether metabolic shifts precede visually apparent rosette formation.

## Acknowledgments

This material is based upon work supported by the National Science Foundation under Grant No. EEC-1648035. The authors gratefully acknowledge assistance by Ying Liu in the Petit Institute for Bioengineering and Biosciences Systems Mass Spectrometry Core. FMF acknowledges support from NIH grants R61CA281667 and R01CA218664.

## Available Code

The analysis package for MALDI rosette characterization is available at https://github.com/kemplab/rosette-radial-gradient

---

**Figure S1:**
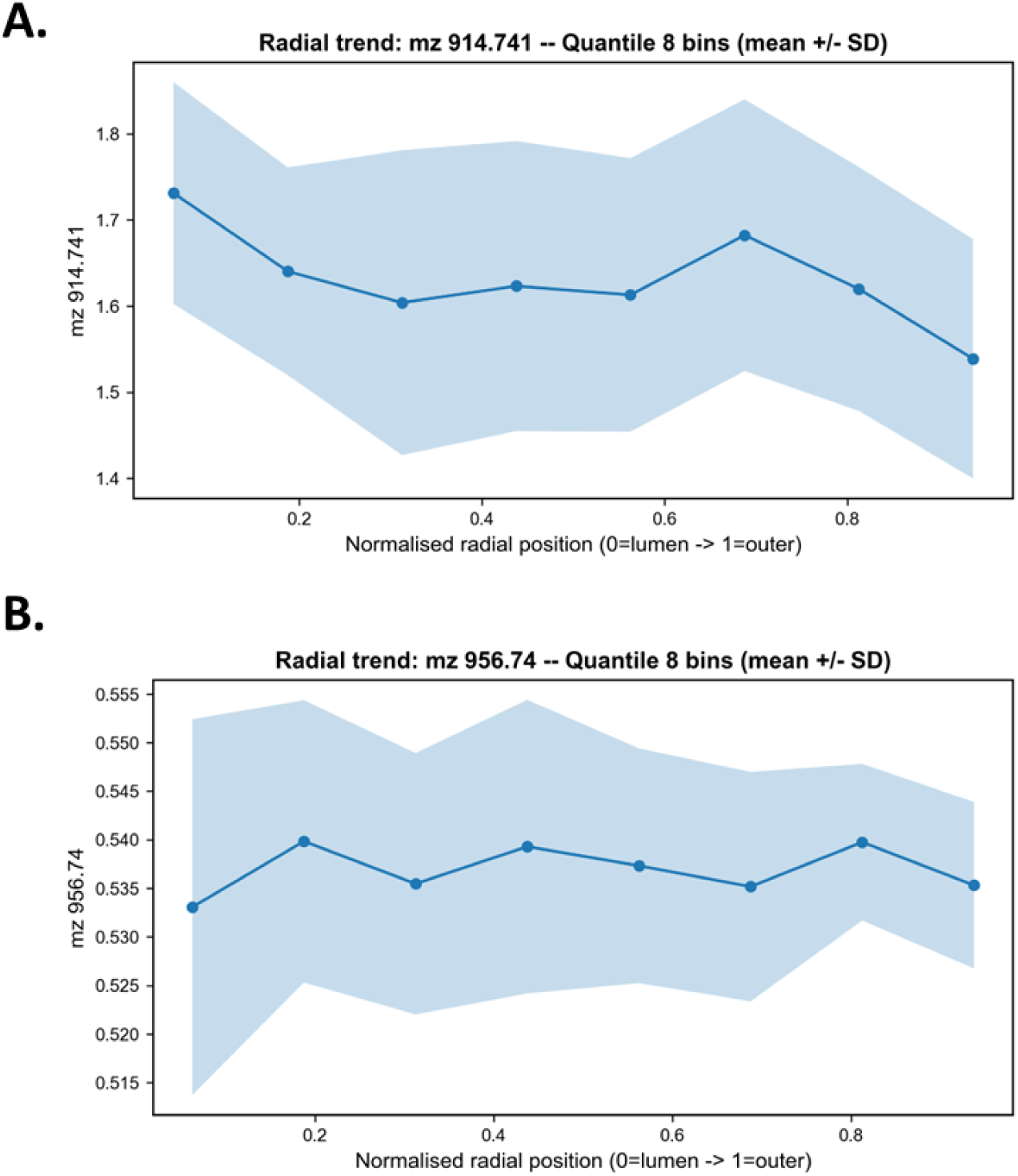
(A) Representative radial intensity profile for a lumen-enriched feature (m/z 914.741; ρ = [-0.548]), showing declining mean intensity from lumen to periphery (mean +/- SD across cells per bin). (B) Representative radial intensity profile for a periphery-associated feature (m/z 956.74; ρ = [0.024]), demonstrating a oscillation with no clear trend. trend.

**Figure S2:**
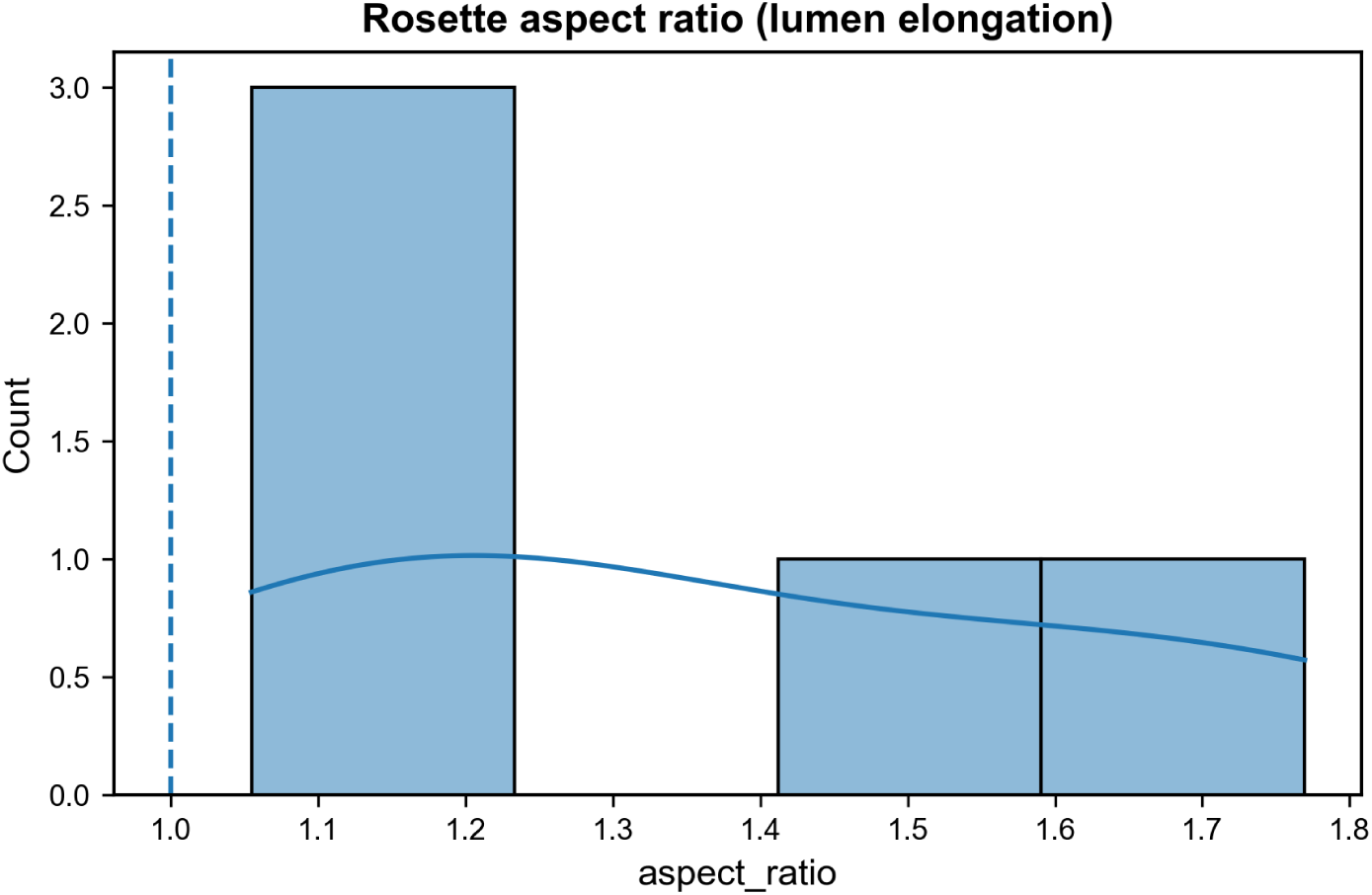
Rosette Circularity Profiling for Cross Rosette Comparison: To quantify rosette shape asymmetry and understand its distribution across neural rosettes detected, principal component analysis (PCA) was applied to the two-dimensional coordinates of Lumen cells within each rosette. The aspect ratio was computed as the ratio of the first to second singular values (σ₁/σ₂), where values exceeding 1.0 indicate elongation along the principal axis. This metric provided quantitative justification for employing the distance-to-boundary model over simpler radial approaches that assume circular symmetry

**Table S1:**
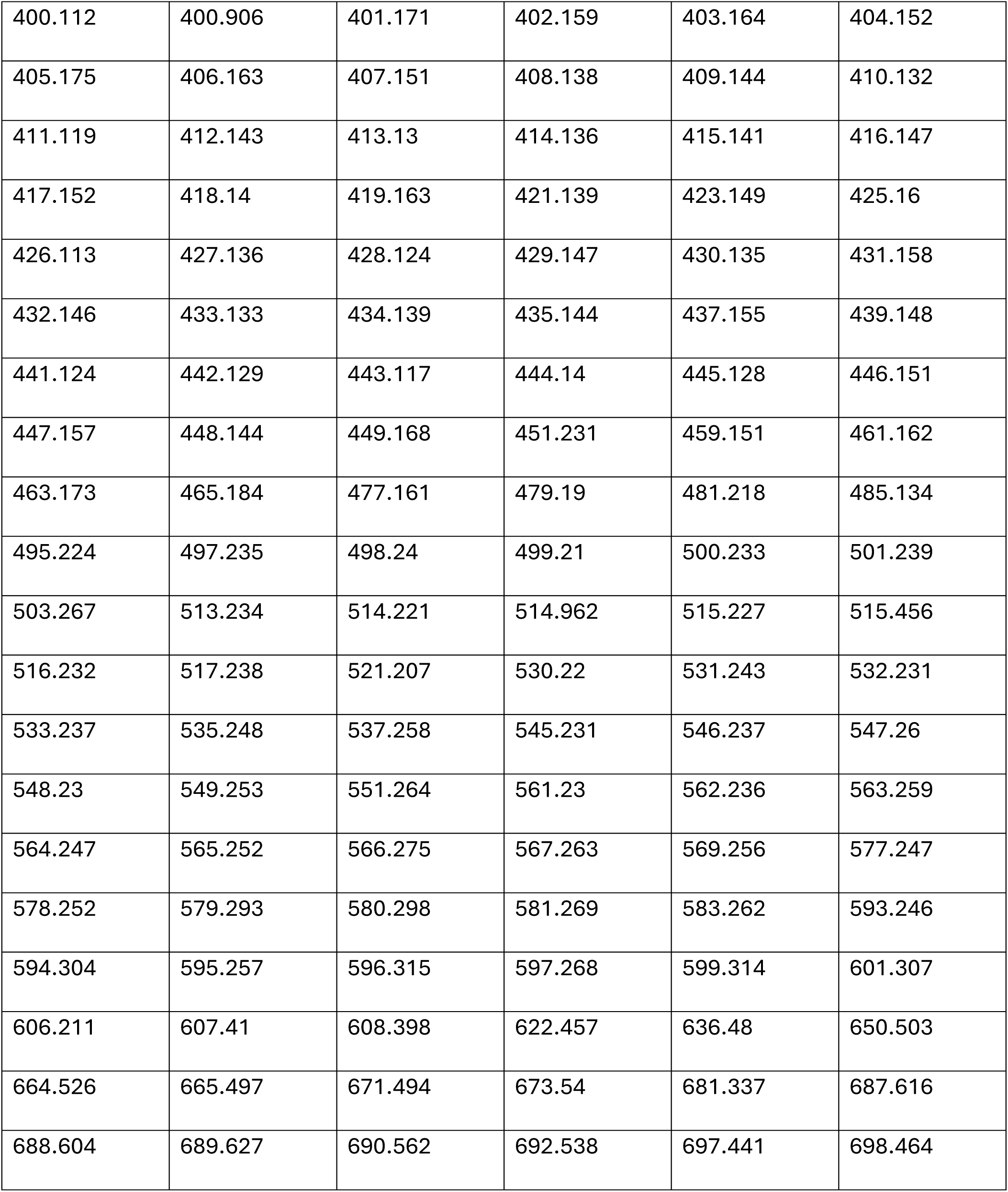
Full list of detected m/z features prior to isotopic peak removal.

**Table S1:**
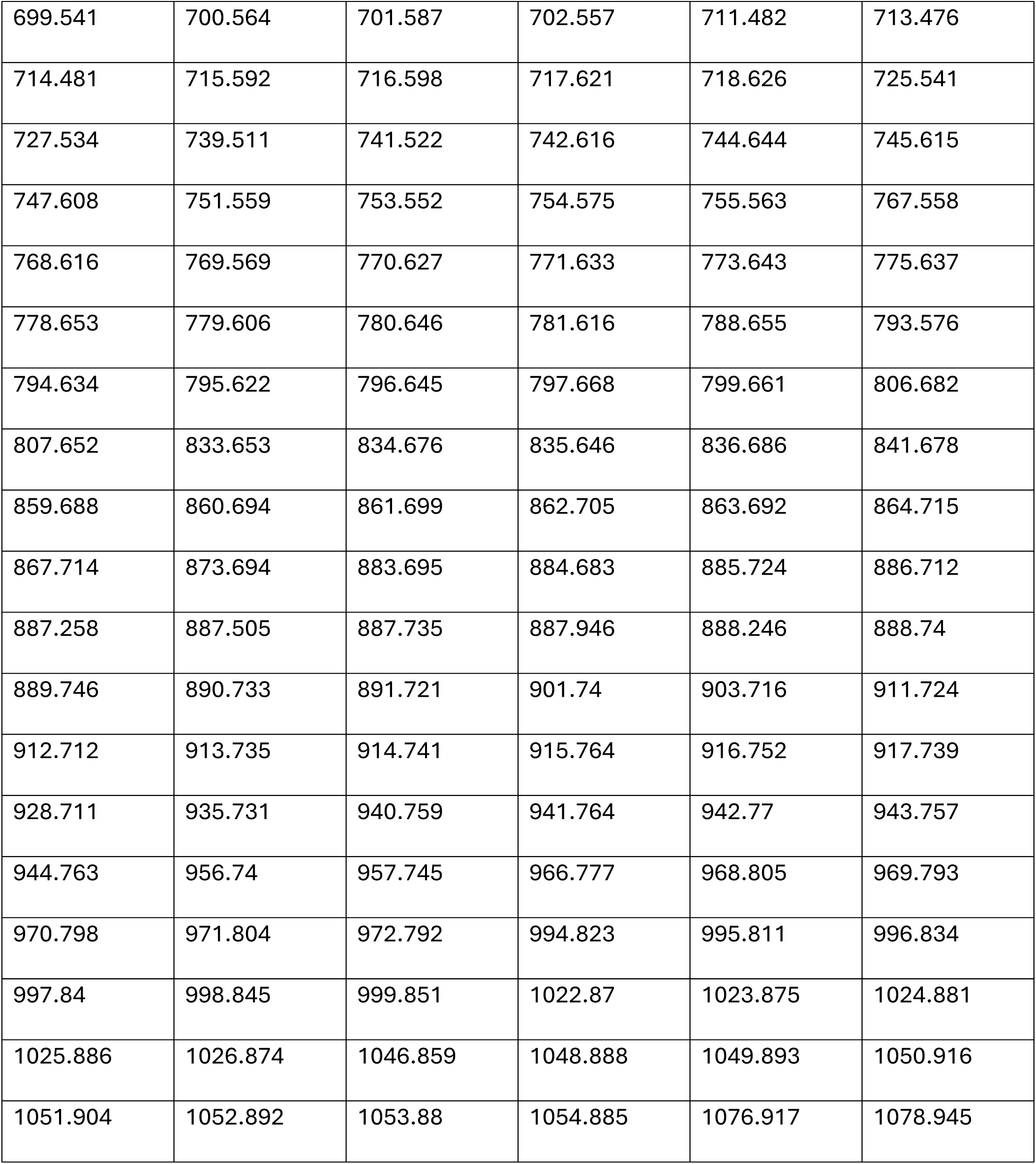
This table includes the complete peak list exported from the raw dataset before any isotope filtering or adduct grouping was applied.

## References

1. Aghayee, A., & Ashton, R. (2021). Methods for Controlled Induction of Singular Rosette Cytoarchitecture Within Human Pluripotent Stem Cell-Derived Neural Multicellular Assemblies. Methods Mol Biol, 2258, 193–203. 10.1007/978-1-0716-1174-6_13

2. Elkabetz, Y., Panagiotakos, G., Al Shamy, G., Socci, N. D., Tabar, V., & Studer, L. (2008). Human ES cell-derived neural rosettes reveal a functionally distinct early neural stem cell stage. Genes Dev, 22(2), 152–165. 10.1101/gad.1616208

3. Filan, C., Charles, S., Casteleiro Costa, P., Niu, W., Cheng, B. F., Wen, Z., Lu, H., & Robles, F. E. (2024). Non-Invasive Label-free Analysis Pipeline for In Situ Characterization of Differentiation in 3D Brain Organoid Models. Res Sq. 10.21203/rs.3.rs-4049577/v1

4. Knight, G. T., Lundin, B. F., Iyer, N., Ashton, L. M., Sethares, W. A., Willett, R. M., & Ashton, R. S. (2018). Engineering induction of singular neural rosette emergence within hPSC-derived tissues. Elife, 7. 10.7554/eLife.37549

5. Lundin, B. F., Knight, G. T., Fedorchak, N. J., Krucki, K., Iyer, N., Maher, J. E., Izban, N. R., Roberts, A., Cicero, M. R., Robinson, J. F., Iskandar, B. J., Willett, R., & Ashton, R. S. (2024). RosetteArray((R)) Platform for Quantitative High-Throughput Screening of Human Neurodevelopmental Risk. bioRxiv. 10.1101/2024.04.01.587605

6. Nikitina, A., Huang, D., Li, L., Peterman, N., Cleavenger, S. E., Fernandez, F. M., & Kemp, M. L. (2020). A Co-registration Pipeline for Multimodal MALDI and Confocal Imaging Analysis of Stem Cell Colonies. J Am Soc Mass Spectrom, 31(4), 986–989. 10.1021/jasms.9b00094

7. Nikitina, A. A., Van Grouw, A., Roysam, T., Huang, D., Fernandez, F. M., & Kemp, M. L. (2023). Mass Spectrometry Imaging Reveals Early Metabolic Priming of Cell Lineage in Differentiating Human-Induced Pluripotent Stem Cells. Anal Chem, 95(11), 4880–4888. 10.1021/acs.analchem.2c04416

8. Priyadarshani, P., Van Grouw, A., Liversage, A. R., Rui, K., Nikitina, A., Tehrani, K. F., Aggarwal, B., Stice, S. L., Sinha, S., Kemp, M. L., Fernandez, F. M., & Mortensen, L. J. (2024). Investigation of MSC potency metrics via integration of imaging modalities with lipidomic characterization. Cell Rep, 43(8), 114579. 10.1016/j.celrep.2024.114579

9. Wilson, P. G., & Stice, S. S. (2006). Development and differentiation of neural rosettes derived from human embryonic stem cells. Stem Cell Rev, 2(1), 67–77. 10.1007/s12015-006-0011-1

10. Ziv, O., Zaritsky, A., Yaffe, Y., Mutukula, N., Edri, R., & Elkabetz, Y. (2015). Quantitative Live Imaging of Human Embryonic Stem Cell Derived Neural Rosettes Reveals Structure-Function Dynamics Coupled to Cortical Development. PLoS Comput Biol, 11(10), e1004453. 10.1371/journal.pcbi.1004453

