## Supplemental Figures for "A Multimodal Workflow for Spatial Metabolic Neighborhood Mapping in Neural Rosette Cultures"

**Supplemental Figures & Tables**

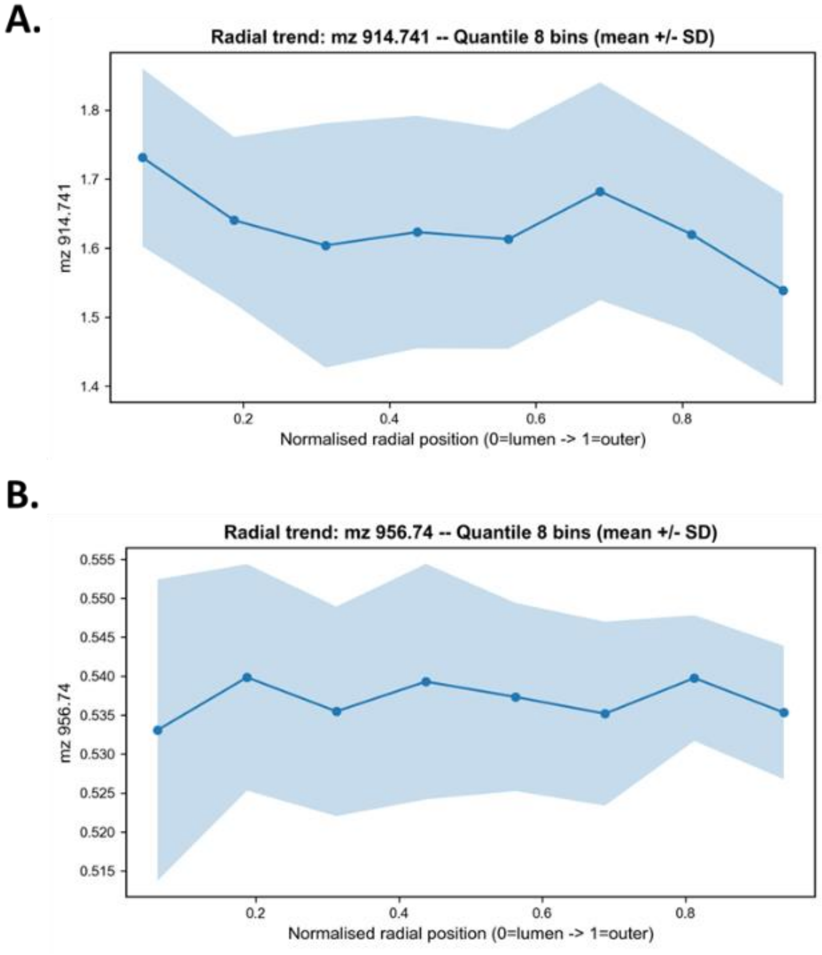


**Figure S1**: (A) Representative radial intensity profile for a lumen-enriched feature (m/z 914.741; ρ = [-0.548]), showing declining mean intensity from lumen to periphery (mean +/- SD across cells per bin). (B) Representative radial intensity profile for a periphery-associated feature (m/z 956.74; ρ = [0.024]), demonstrating a oscillation with no clear trend.

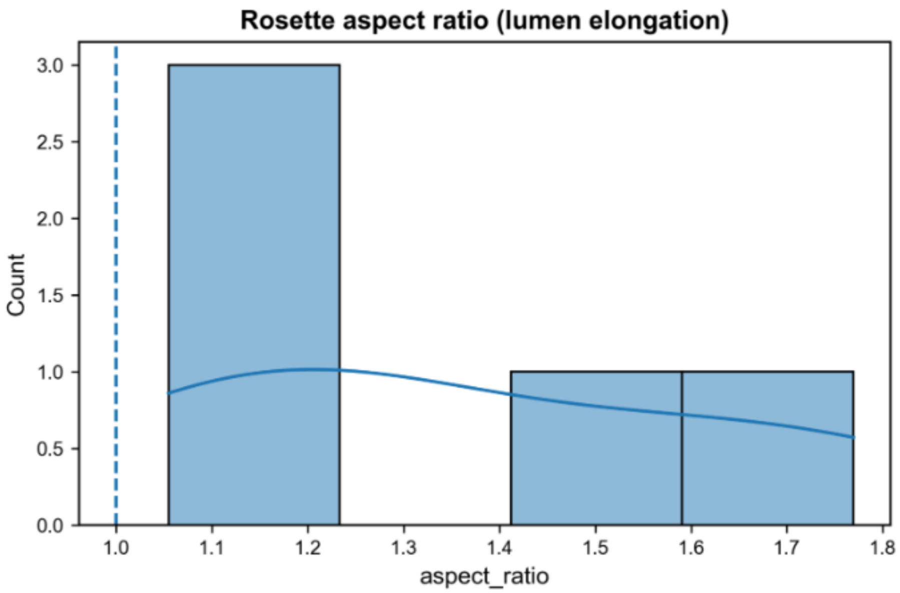


**Figure S2:**

**Rosette Circularity Profiling for Cross Rosette Comparison:** To quantify rosette shape asymmetry and understand its distribution across neural rosettes detected, principal component analysis (PCA) was applied to the two-dimensional coordinates of Lumen cells within each rosette. The aspect ratio was computed as the ratio of the first to second singular values ($\sigma_1/\sigma_2$), where values exceeding 1.0 indicate elongation along the principal axis. This metric provided quantitative justification for employing the distance-to-boundary model over simpler radial approaches that assume circular symmetry.

**Table S1: Full list of detected m/z features prior to isotopic peak removal.**

| | | | | | |
|---|---|---|---|---|---|
| 400.112 | 400.906 | 401.171 | 402.159 | 403.164 | 404.152 |
| 405.175 | 406.163 | 407.151 | 408.138 | 409.144 | 410.132 |
| 411.119 | 412.143 | 413.13 | 414.136 | 415.141 | 416.147 |
| 417.152 | 418.14 | 419.163 | 421.139 | 423.149 | 425.16 |
| 426.113 | 427.136 | 428.124 | 429.147 | 430.135 | 431.158 |
| 432.146 | 433.133 | 434.139 | 435.144 | 437.155 | 439.148 |
| 441.124 | 442.129 | 443.117 | 444.14 | 445.128 | 446.151 |
| 447.157 | 448.144 | 449.168 | 451.231 | 459.151 | 461.162 |
| 463.173 | 465.184 | 477.161 | 479.19 | 481.218 | 485.134 |
| 495.224 | 497.235 | 498.24 | 499.21 | 500.233 | 501.239 |
| 503.267 | 513.234 | 514.221 | 514.962 | 515.227 | 515.456 |
| 516.232 | 517.238 | 521.207 | 530.22 | 531.243 | 532.231 |
| 533.237 | 535.248 | 537.258 | 545.231 | 546.237 | 547.26 |
| 548.23 | 549.253 | 551.264 | 561.23 | 562.236 | 563.259 |
| 564.247 | 565.252 | 566.275 | 567.263 | 569.256 | 577.247 |
| 578.252 | 579.293 | 580.298 | 581.269 | 583.262 | 593.246 |
| 594.304 | 595.257 | 596.315 | 597.268 | 599.314 | 601.307 |
| 606.211 | 607.41 | 608.398 | 622.457 | 636.48 | 650.503 |
| 664.526 | 665.497 | 671.494 | 673.54 | 681.337 | 687.616 |

| | | | | | |
|---|---|---|---|---|---|
| 688.604 | 689.627 | 690.562 | 692.538 | 697.441 | 698.464 |
| 699.541 | 700.564 | 701.587 | 702.557 | 711.482 | 713.476 |
| 714.481 | 715.592 | 716.598 | 717.621 | 718.626 | 725.541 |
| 727.534 | 739.511 | 741.522 | 742.616 | 744.644 | 745.615 |
| 747.608 | 751.559 | 753.552 | 754.575 | 755.563 | 767.558 |
| 768.616 | 769.569 | 770.627 | 771.633 | 773.643 | 775.637 |
| 778.653 | 779.606 | 780.646 | 781.616 | 788.655 | 793.576 |
| 794.634 | 795.622 | 796.645 | 797.668 | 799.661 | 806.682 |
| 807.652 | 833.653 | 834.676 | 835.646 | 836.686 | 841.678 |
| 859.688 | 860.694 | 861.699 | 862.705 | 863.692 | 864.715 |
| 867.714 | 873.694 | 883.695 | 884.683 | 885.724 | 886.712 |
| 887.258 | 887.505 | 887.735 | 887.946 | 888.246 | 888.74 |
| 889.746 | 890.733 | 891.721 | 901.74 | 903.716 | 911.724 |
| 912.712 | 913.735 | 914.741 | 915.764 | 916.752 | 917.739 |
| 928.711 | 935.731 | 940.759 | 941.764 | 942.77 | 943.757 |
| 944.763 | 956.74 | 957.745 | 966.777 | 968.805 | 969.793 |
| 970.798 | 971.804 | 972.792 | 994.823 | 995.811 | 996.834 |
| 997.84 | 998.845 | 999.851 | 1022.87 | 1023.875 | 1024.881 |
| 1025.886 | 1026.874 | 1046.859 | 1048.888 | 1049.893 | 1050.916 |
| 1051.904 | 1052.892 | 1053.88 | 1054.885 | 1076.917 | 1078.945 |

**Table S1:** This table includes the complete peak list exported from the raw dataset before any isotope filtering or adduct grouping was applied.
